# Molecular characterisation of common *Culicoides* Biting Midges (Diptera: Ceratopogonidae) in Ireland

**DOI:** 10.1101/2025.01.08.631704

**Authors:** Elsie Isiye, Angela Valcarcel Olmeda, Thomas Curran, David O’Neill, Theo de Waal, Gerald Barry, Aidan O’Hanlon, James O’Shaughnessy, Nicole Keohane McCarthy, Akke Vellinga, Audrey Jenkinson, Alan Johnson, Damien Barrett, Sarah Costello, Annetta Zintl, Denise O’Meara

**Author notes:** Joint last authors (Annetta Zintl, Denise O’Meara).

## Abstract

**Background:** Biting midges in the genus *Culicoides* (Diptera: Ceratopogonidae) act as vectors for several arboviruses, including Bluetongue virus (BTV) and Schmallenberg virus (SBV), which affect livestock health and productivity. In Ireland, limited genetic data are available regarding the diversity of *Culicoides* species. This study represents the first attempt to characterise *Culicoides* in this region using molecular techniques.

**Methods:** Adult *Culicoides* samples were captured using Onderstepoort Veterinary Institute (OVI) traps across six locations in Ireland. Subsequent molecular analyses involved polymerase chain reaction (PCR) and sequencing of the cytochrome oxidase subunit 1 (CO1) and the internal transcriber spacer (ITS) barcoding regions to obtain species identities. Additionally, using both markers, we inferred the population genetic structure and potential colonisation pathways of *Culicoides obsoletus sensu stricto (s. str.)*, the major vector species in Ireland.

**Results:** DNA barcoding facilitated identification of 177 specimens. Eight common *Culicoides* species were identified through DNA barcoding of CO1 and ITS gene regions. The presence of putative vectors of Bluetongue virus (BTV) and Schmallenberg virus (SBV) were also confirmed, including species in the subgenus *Avaritia* (*C. obsoletus s. str., C. scoticus, C. chiopterus, and C. dewulfi*) and subgenus *Culicoides s. str.* (*C. pulicaris* and *C. punctatus*). Phylogenetic analysis confirmed the relationship between these vector species and facilitated the placement of *Culicoides* sp. which could not be identified to species level through DNA barcoding. Haplotype network analysis of *C. obsoletus* showed that some haplotypes of these species are shared between Continental Europe, the UK, and Ireland, suggesting a possible incursion pathway for this vector.

**Conclusion:** DNA barcoding employing a combination of two barcodes, CO1 and ITS, proved effective in identifying *Culicoides*, especially species within the *obsoletus* complex, which are difficult to morphologically distinguish. Our findings also suggest that investigation of the population genetic structure of *Culicoides* spp. could be used to model the potential introduction routes of midge-borne pathogens into the country.

## Introduction

There are over 1,400 species of biting midges in the genus *Culicoides* Latreille, 1809 (Diptera: Ceratopogonidae) and they are found worldwide in all landmasses except Antarctica [1]. Their broad distribution is attributed to various factors, such as the availability of different ecological niches, breeding sites, and hosts [2]. Identification of biting midges has become increasingly important because they play a central role in the transmission of several pathogens to humans and animals, including arboviruses, bacteria, protozoa, and parasitic nematodes [3, 4]. In Europe, they are known to transmit Bluetongue virus (BTV), Schmallenberg virus (SBV), Epizootic Haemorrhagic Disease Virus (EHDV) (5), and African Horse Sickness virus (AHSV) [1, 6, 7].

BTV and SBV have significant economic and animal health implications in Europe. In the early 2000s, the number of BTV cases in Europe increased dramatically, with a particularly severe outbreak that occurred between 2006 and 2008. During this period, BTV serotype 8 (BTV-8) caused economic losses owing to decreased production, mortality, and costs related to control and management measures [8, 9]. In the case of SBV, following its discovery in Germany in 2011, it spread across Europe, causing fever, reduced milk yield, abortions, stillbirths and congenital abnormalities in ruminants [10–13]. While BTV is absent from Ireland, SBV was first detected in 2012 causing a major outbreak in 2016 and 2017 and is now considered endemic in Ireland [14–17].

Following the BTV-8 outbreak from 2006 to 2008 and the SBV outbreak from 2011 to 2012 in Northern Europe, surveillance programmes were established as part of contingency plans to control arboviruses transmitted by biting midges. Various species within different subgenera, including *Culicoides chiopterus* Meigen, 1830, *Culicoides dewulfi* Goetghebuer, 1936, *Culicoides obsoletus* Meigen, 1818, *Culicoides scoticus* Downes & Kettle, 1952 (subgenus *Avaritia* Fox, 1955), *Culicoides lupicaris* Downes & Kettle, 1952, *Culicoides punctatus* Meigen, 1804, and *Culicoides pulicaris* Linnaeus, 1758 (subgenus *Culicoides s. str.*), have been identified as potential vectors of BTV [18, 19].

In Ireland, the Department of Agriculture, Food, and the Marine (DAFM) carried out the first surveillance of *Culicoides* from 2007 to 2009 as part of the National BTV Vector Surveillance Programme. This study was prompted by the outbreak of BTV in Europe and aimed to identify potential vector species. Several suspected vector species were identified, with the most abundant species including *C. obsoletus sensu lato* (*s. lat.*), *C. dewulfi*, *C. chiopterus*, *C. pulicaris*, and *C. punctatus* collectively accounting for 80–90% of all identified *Culicoides* [20].

In response to the 2012/2013 SBV epidemic in Ireland [14, 15], a sentinel herd surveillance study was initiated across 26 livestock farms in the south of Ireland to monitor the post-epizootic circulation of the virus [16]. *Culicoides* surveillance was later conducted on ten of these farms, and it was reported that the most abundant species identified were *C. obsoletus/C. scoticus* (38%), *C. dewulfi* (36%), *C. chiopterus* (5%), *C. pulicaris* (9%), and *C. punctatus* (5%), collectively accounting for 93% of all *Culicoides* collected. In this study, 20 species of *Culicoides* biting midges were identified, including one species not previously known from Ireland, *C. cameroni* Campbell and Pelham-Clinton, 1960, which raised Ireland’s species list to 31 species [21], belonging to eight subgenera. As these previous studies predominantly relied on morphological examinations for species identification, there is a lack of genetic information regarding *Culicoides* in Ireland, which has inhibited the development of rapid molecular identification techniques.

Regarding the introduction of SBV into Ireland in 2012, several transmission routes have been suggested, including the importation of infected animals and windborne dispersal of infected midges. Surveillance in 2012 and 2013 identified a high concentration of SBV cases in the southeast of the island [16] suggesting that SBV may have been introduced via winds carrying infected midges from the UK or continental Europe. Moreover, McGrath et al. [22] utilised serological analysis of archived bovine sera to identify potential dispersal windows and atmospheric dispersion modelling (ADM) to evaluate environmental conditions conducive to the transportation of *Culicoides* into Ireland. Atmospheric dispersion modelling pointed to favourable conditions for midge dispersal from southern England in August 2012, though long-range transportation events were rare with only one significant instance identified during the 2012 vector season. These studies and hypotheses highlight the need for further research to reveal potential routes for the spread of midges and midge-borne viruses into Ireland.

Morphological identification of *Culicoides* species is time-consuming and requires highly trained personnel. Moreover, they are difficult to identify cryptic species and can result in misidentifications. DNA barcoding of a common gene region has enabled the accurate and reliable molecular identification and phylogenetic placement of *Culicoides* species [23–25] and has helped to catalogue previously hidden diversity within this taxonomic group in various countries [25–30]. Molecular data have also proven invaluable in population genetics studies, particularly for tracing the colonization history of biting midges. These studies indicate that, while native populations exhibit strong genetic structuring, recently colonized regions often show low genetic differentiation. It has been suggested that this pattern which has been observed in important vector species such as *C. obsoletus sensu lato*, *C. imicola* Kieffer, 1913, *C. brevitarsis* Kieffer, 1917 and *C. mahasarakhamense* could potentially be used to investigate the incursion and spread of pathogens that they transmit [31–35].

The present study aimed to conduct the first comprehensive molecular characterisation of common biting midge species found in Ireland. In addition, we undertook a population genetic analysis of the vector species, *C. obsoletus s. str.*, to determine the genetic diversity and relationship of Irish specimens to populations characterised elsewhere, potentially revealing migration patterns and sources of incursion.

## Materials and methods

### Sample collection

Onderstepoort Veterinary Institute (OVI) ultraviolet (UV) light suction traps were used to collect adult insect samples as part of the Network of Insect Vectors (NetVec) Ireland project https://www.ucd.ie/netvecireland/. Each trap was operated overnight (from dusk to dawn), and samples were collected in beakers, transferred into 70% ethanol, and stored at room temperature until specimen identification. Samples were pre-identified morphologically to the complex/group level and, where possible, to the species level using interactive keys described by Mathieu et al. [36]. A subset of samples from the six study sites (Figure 1) collected between June and July 2022 were then used for characterization using molecular methods.

**Figure 1:**
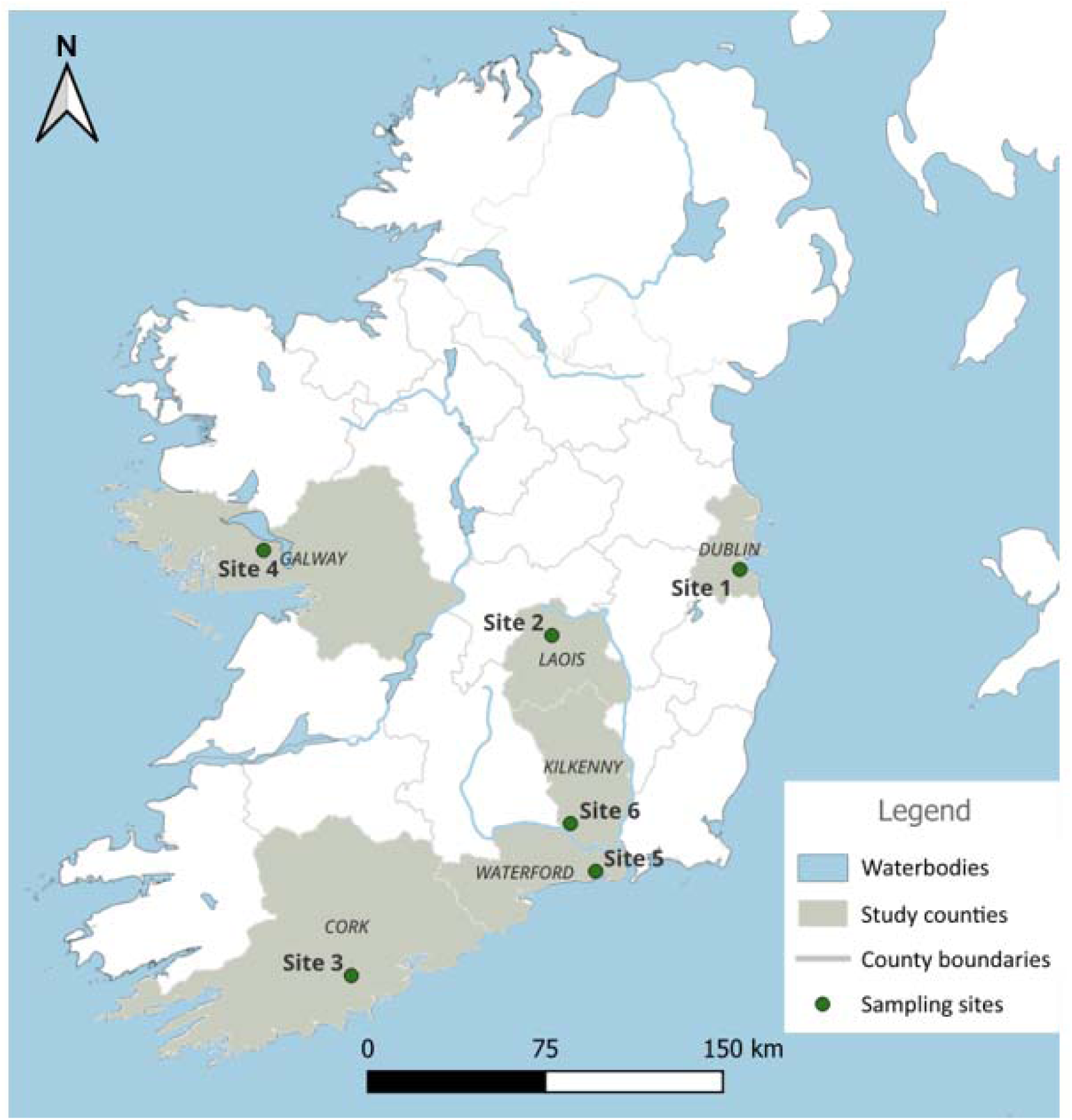
Location of sample collection sites in six counties in Ireland during the midge surveillance activities conducted in June and July 2022. The map was generated in QGIS 3.38.1 with shape files provided by Natural Earth (http://www.naturalearthdata.com/)

### DNA extraction, amplification and sequencing

Whole individual biting midges were air-dried for one hour to remove ethanol and subsequently crushed individually in 1.5 mL microcentrifuge tubes. DNA was extracted from each crushed midge using the NucleoSpin DNA Extraction Kit (Macherey-Nagel), following the manufacturer’s protocol and eluted in 100 µL of elution buffer. The purity and concentration of the extracted DNA were measured using a NanoDrop™ 8000 Spectrophotometer (Thermo Scientific).

*Culicoides* DNA samples were amplified in the CO1 region with primers LCO1490/HCO2198 primers [37] as well as ITS regions using the ITSA/ITS2B primers [38] and PanCul [6]. PCR reactions were performed in a final volume of 10 μl consisting of 5 µL GoTaq™ Hot Start Polymerase: Green Master Mix, 2X (Promega), 3 µL nuclease-free water, 1 µl of a primer mix containing 5 µm each of the forward and reverse primers and 1 µL of the template. Negative controls contained 1 µl of nuclease-free water in lieu of template DNA. The PCR cycling conditions used for CO1 included an initial denaturation at 95°C for 15 min, followed by 40 cycles of 95 °C for 40 s, 46 °C for 40 s, 72 °C for 40 s, and a final extension at 72 °C for 7 min a final hold at 4 °C. For ITS 2 and PanCul the PCR cycling conditions were as follows; an initial denaturation at 95 °C for 3 min followed by 35 cycles of 95 °C for 1 min, 54 °C for 1min, 72 °C for 1 min, and a final extension at 72 °C for 20 min and a final hold at 4 °C. Following the amplification of DNA, agarose gel electrophoresis was used to visualise the PCR products on a 1.6% agarose gel. PCR products of the expected size were purified using microCLEAN (Microzone) following the manufacturer’s protocol and sequenced in the forward direction on an Applied Biosystems 3500 Genetic Analyser using the BigDye Terminator Cycle Sequencing Kit V3.1 (Applied Biosystems).

The quality of sequences generated was evaluated on Chromas (Version: 2.6.6) before performing the standard nucleotide Basic Local Alignment Search Tool (BLASTn) on GenBank. Sample identity was verified based on DNA sequence similarity with reference sequences from the GenBank public repository, using default search parameters. BLAST hits with the highest query cover score and percentage identity values were considered candidate species (Additional file 1).

For specimens with lower sequencing quality, amplicons were cloned into the pDrive Cloning Vector and transformed into competent Top Ten cells using the Qiagen PCR Cloning Kit, following the manufacturer’s protocol, as described in previous studies. [30, 39, 40]. Direct colony PCR was performed to confirm the presence of inserts in transformants using M13 primers (Table 1). The PCR reaction mixture for each reaction comprised 5 µL GoTaq™ Hot Start Polymerase: Green Master Mix, 2X (Promega), 3 µL nuclease-free water, 5 µm of the forward and reverse M13 primers, and 1 µL template. Negative controls contained 1 µl of nuclease-free water in lieu of template DNA. The PCR conditions included an initial step at 95 °C for 10 min, followed by 30 cycles at 95 °C for 30 s, 57 °C for 30 s, 72 °C for 1 min and 72 °C for 10 min. One microlitre of each PCR product was visualised using 1% agarose gel electrophoresis. Positive PCR products were purified and sequenced as previously described.

**Table 1:**
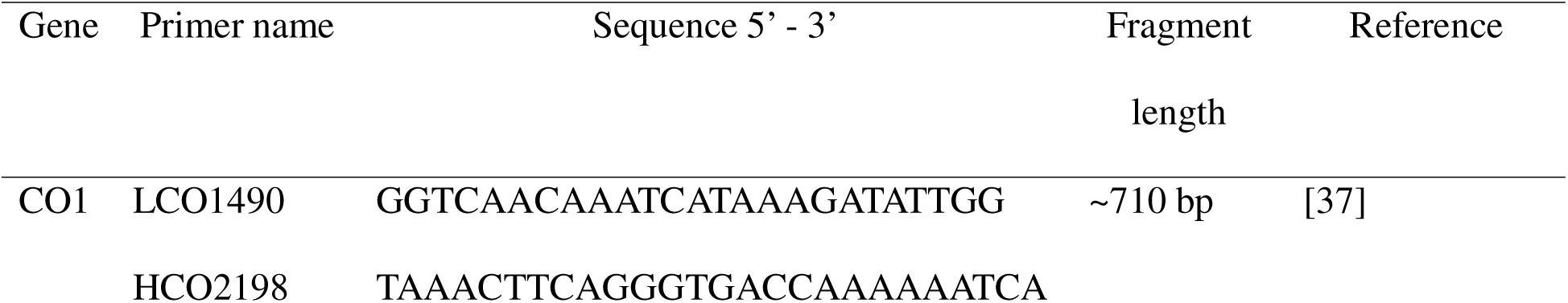
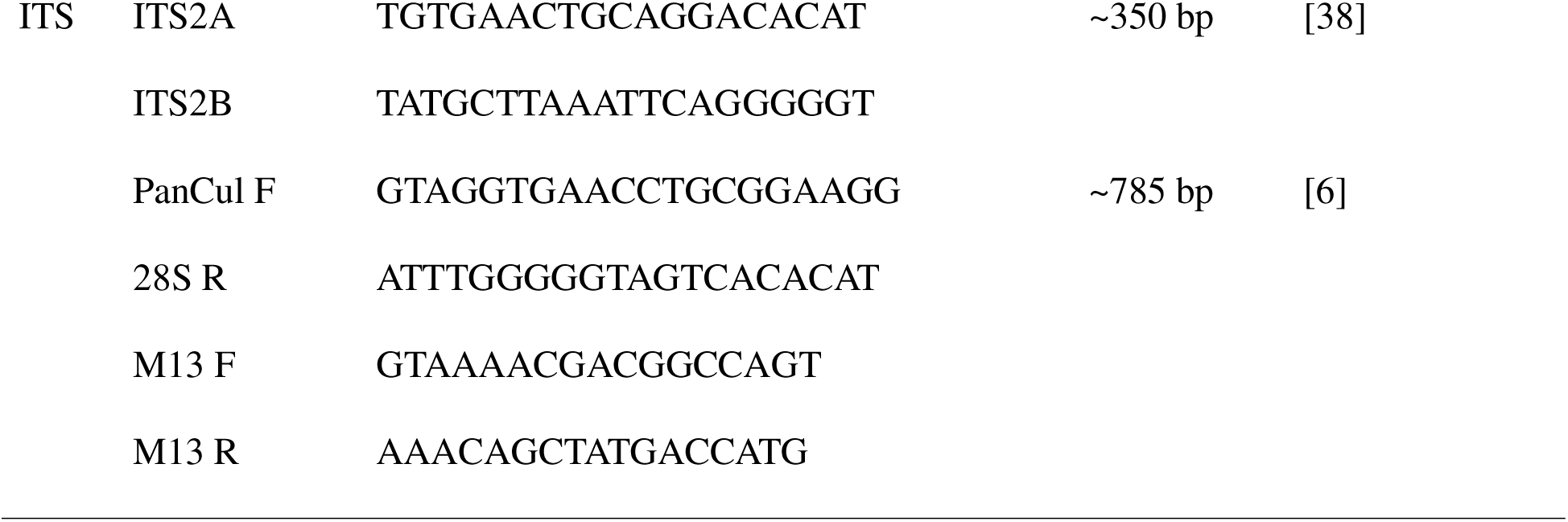
Primers used in DNA amplification and sequencing.

### Culicoides species identification

To visualize the diversity of common *Culicoides* species in the subgenus *Avaritia* and *Culicoides* trapped in the six study sites, a chord diagram was constructed using the “circlize” package in RStudio [41]. The chord diagram illustrates the interconnections and diversity of *Culicoides* species between various sites.

### Phylogenetic analysis

To examine the genetic relationships among species in this study, multiple sequence alignments of CO1 and the ITS sequences were performed using the online versions of MAFFT [42] and Molecular Evolutionary Genetics Analysis (MEGA) software version 11 [43]. In addition to the sequences generated in this study, CO1 and ITS sequences of the same species obtained from GenBank (Additional File 2) were incorporated into the alignments to facilitate a broader comparison. The CO1 alignment was truncated to 507 base pairs (bp), while the ITS alignment was trimmed to 218 bp, with gaps excluded from the ITS alignment using MEGA. The ITS dataset included a total of 69 sequences, encompassing both newly generated and previously published sequences and similarly, the CO1 dataset comprised 59 sequences. Neighbour-joining phylogenetic trees based on both the ITS and CO1 sequences were constructed, incorporating reference sequences obtained from GenBank. To assess the robustness of the phylogenetic trees, the bootstrapping method with 1,000 bootstrap replications was utilised. The resulting trees were visualized using the online tool Interactive Tree of Life (iTOL, available at https://itol.embl.de/).

### Haplotype analysis of Culicoides obsoletus

In this study, 53 CO1 and 62 ITS gene sequences of *Culicoides obsoletus* were aligned using the CLUSTAL W method in MEGA. These sequences were compared with reference sequences from GenBank, representing different *C. obsoletus* haplotypes (Additional file 3).

For the CO1 alignment, reference sequences were sourced from *C. obsoletus* populations in the UK (n=9) as well as from multiple countries across Continental Europe and neighbouring regions, including France (n=2), Switzerland (n=2), Italy (n=2), Germany (n=3), the Netherlands (n=3), Spain (n=3), Norway (n=3), Greece (n=1), Denmark (n=3), Poland (n=3), Serbia (n=3), Bulgaria (n=3), Macedonia (n=3), Morocco (n=4) and Turkey (n=2). For the ITS locus fewer reference sequences were available and the following sequences were included: UK (n=9), Italy (n=15), France (n=5), Norway, and Germany (n=3). A total of 102 CO1 and 86 ITS2 sequences were truncated to 528 bp and 246 bp, respectively. Both alignments were exported as NEXUS files into Population Analysis with Reticulate Trees (PopART; http://popart.otago.ac.nz/), and a Median-Joining Network diagram illustrating haplotype diversity, was constructed using the median algorithm with default settings [44].

## Results

### Genetic identification of common Culicoides *spp*

Overall, 177 specimens were identified by simultaneous sequencing of the CO1 and ITS sequences. Both barcodes were successful in determining species identities, with the sequencing results from each barcode region complementing the other. Of the 177 specimens, 139 (78.5%) were identified through their CO1 region, and 166 (93.2%) were identified by the ITS region. Overall, eight species were identified. *Culicoides obsoletus* represented the most dominant species in the sample set with 34.4% of specimens identified as the species (62 flies), followed by *C. impunctatus* at 20.6% (36 flies), *C. scoticus* at 16.1% (27 flies), *C. chiopterus* at 11.1% (20 flies), and *C. dewulfi* at 9.4% (17 flies). The least common species were *C. pulicaris* (3.9%, 7 flies), *C. punctatus* (2.8%, 5 flies), and an unidentified *Culicoides* sp. (1.7%, 3 flies). In most cases, the ITS2 barcode was more effective, particularly in identifying the closely related species *C. scoticus* and *C. obsoletus s. str.* within the *obsoletus* complex.

The chord diagram (Fig. 2) shows the distribution of the eight species across the six sampling sites. *Culicoides obsoletus s. str.* was detected at all six sites while the other species, *C. scoticus*, *C. punctatus*, *C. impunctatus*, *C. dewulfi*, and *C. chiopterus*, were only detected at some sites. Two *Culicoides* samples from site 3 (Laois) (Fig. 2) amplified and sequenced using ITS2 primers only, yielding short sequences of approximately 108 base pairs, which hindered analysis. Following cloning and resequencing, approximately 380 base pair sequences were produced, enabling species identification. Nucleotide BLAST analysis revealed 99% and 98% similarity to an unidentified *Culicoides* spp reported by Gomulski et al. [39] (GenBank accessions AY599813 and AY599814, respectively). A third sample from the same site, displaying similar characteristics, was also identified as *Culicoides* sp. with approximately 98% similarity to the sequence deposited under GenBank accession number AY599815.

**Figure 2:**
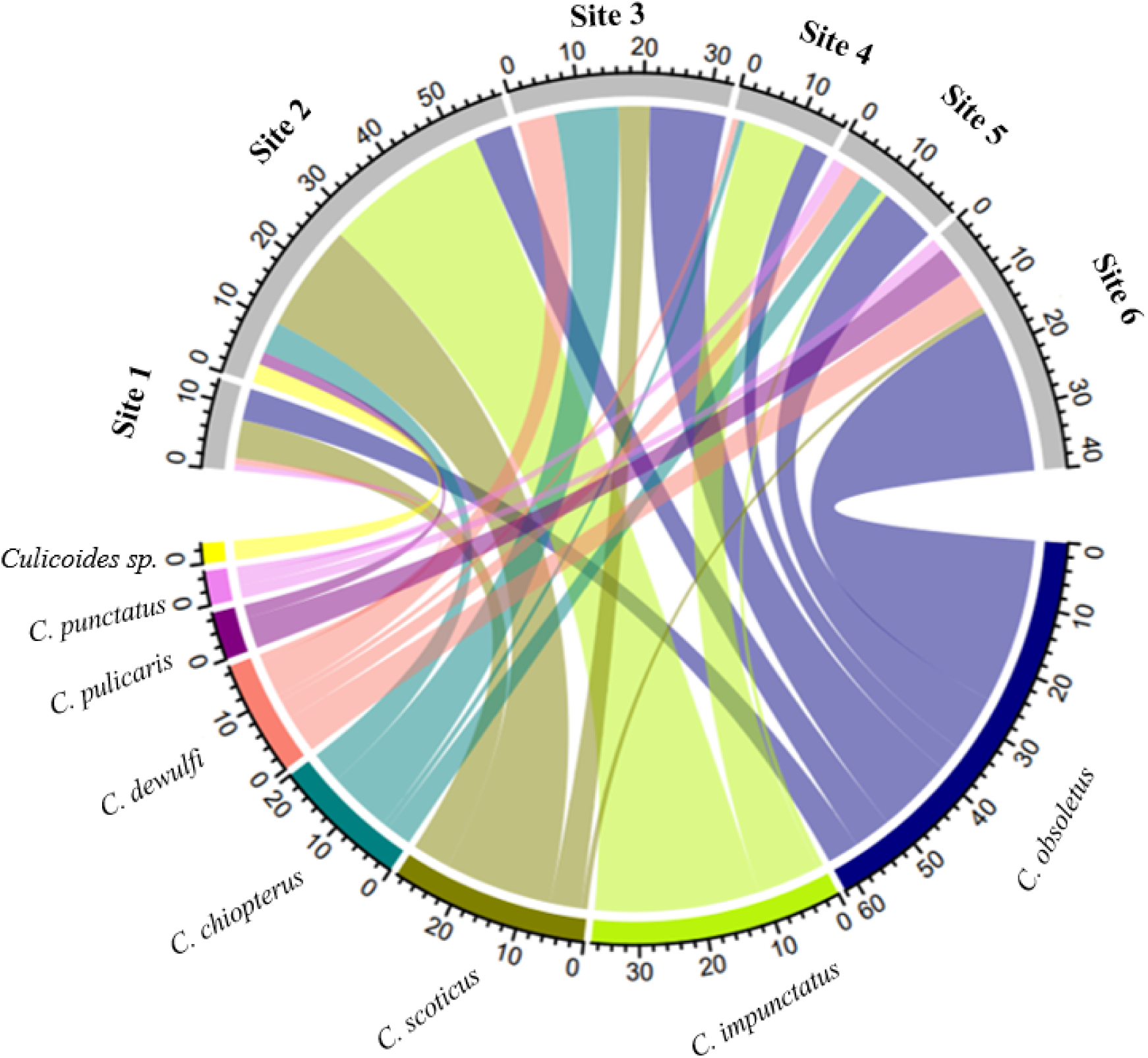
Species diversity across the six sample sites: Sites 1 (Dublin Site), Site 2 (Laois), Site 3 (Cork), Site 4 (Galway), Site 5 (Waterford), and Site 6 (Kilkenny).

Regarding *Culicoides* species diversity in the sampled locations, site 2 (Laois), site 4 (Galway), and site 5 (Waterford) exhibited the highest diversity, with eight *Culicoides* species recorded at each site. Site 3 (Cork) and site 6 (Kilkenny) each had five different species and finally, site 1 (Dublin) showed the lowest diversity, with only four species identified.

### Phylogenetic analysis of the Culicoides species

Phylogenetic trees representing the two gene regions were generated with branches of the tree depicting the evolutionary relationships among *Culicoides* species (Figs. 3 and 4). Based on the CO1 and ITS2 sequences, two subgenera, *Avaritia* and *Culicoides s. str.* were identified. Bootstrap values are shown on a gradient from red to green, indicating the reliability of these inferred relationships, with red representing lower confidence (18/19%) and green representing higher confidence (100%).

**Figure 3:**
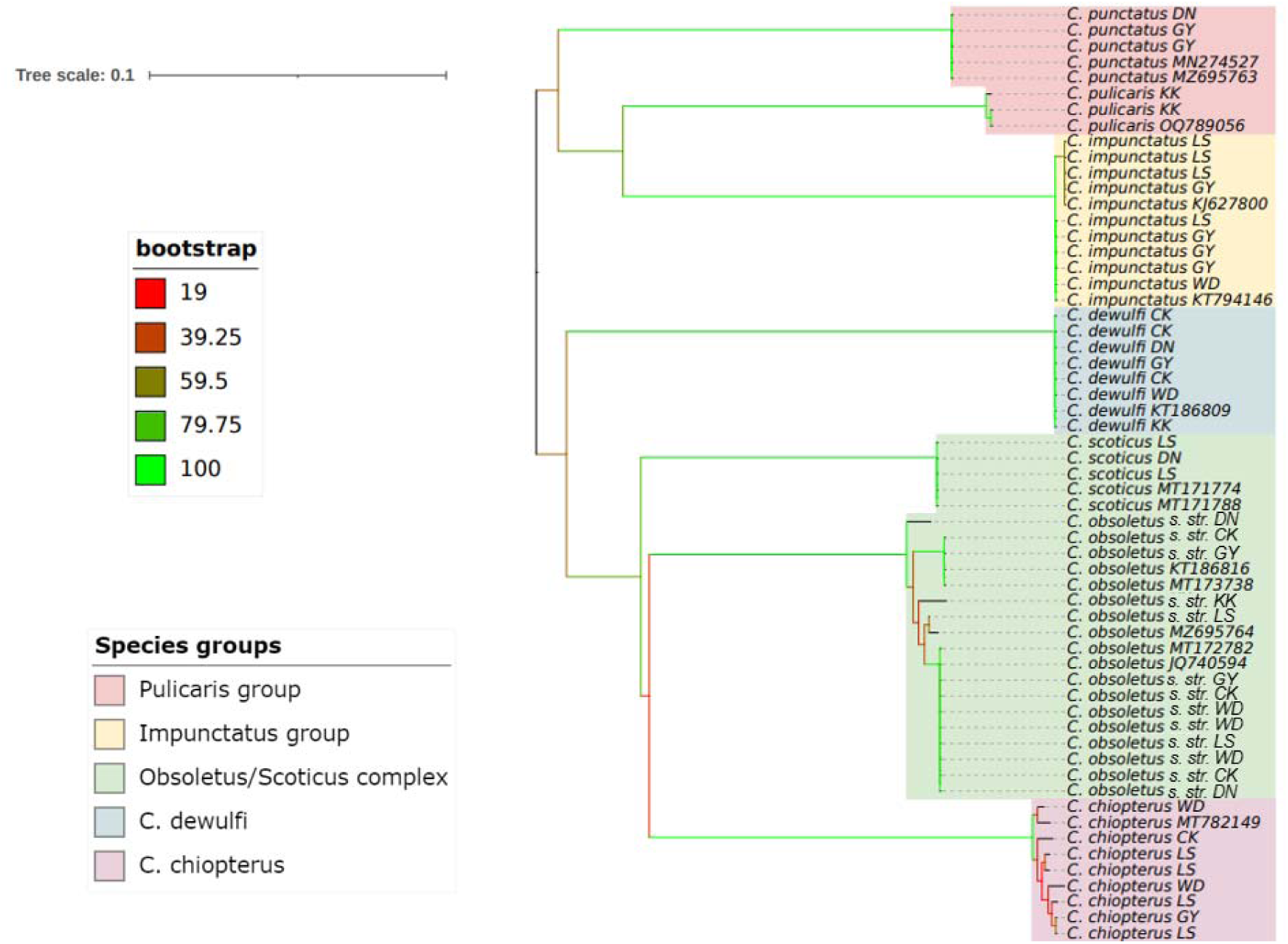
Neighbour-joining phylogenetic tree generated from an alignment of identified Irish CO1 *Culicoides* sequences and reference sequences obtained from GenBank truncated to a final sequence length of 507 bp. Labels on the right side indicate the species name and county code (WD – Waterford; KK – Kilkenny; CK – Cork; DN – Dublin; LS – Laois and GY – Galway).

**Figure 4:**
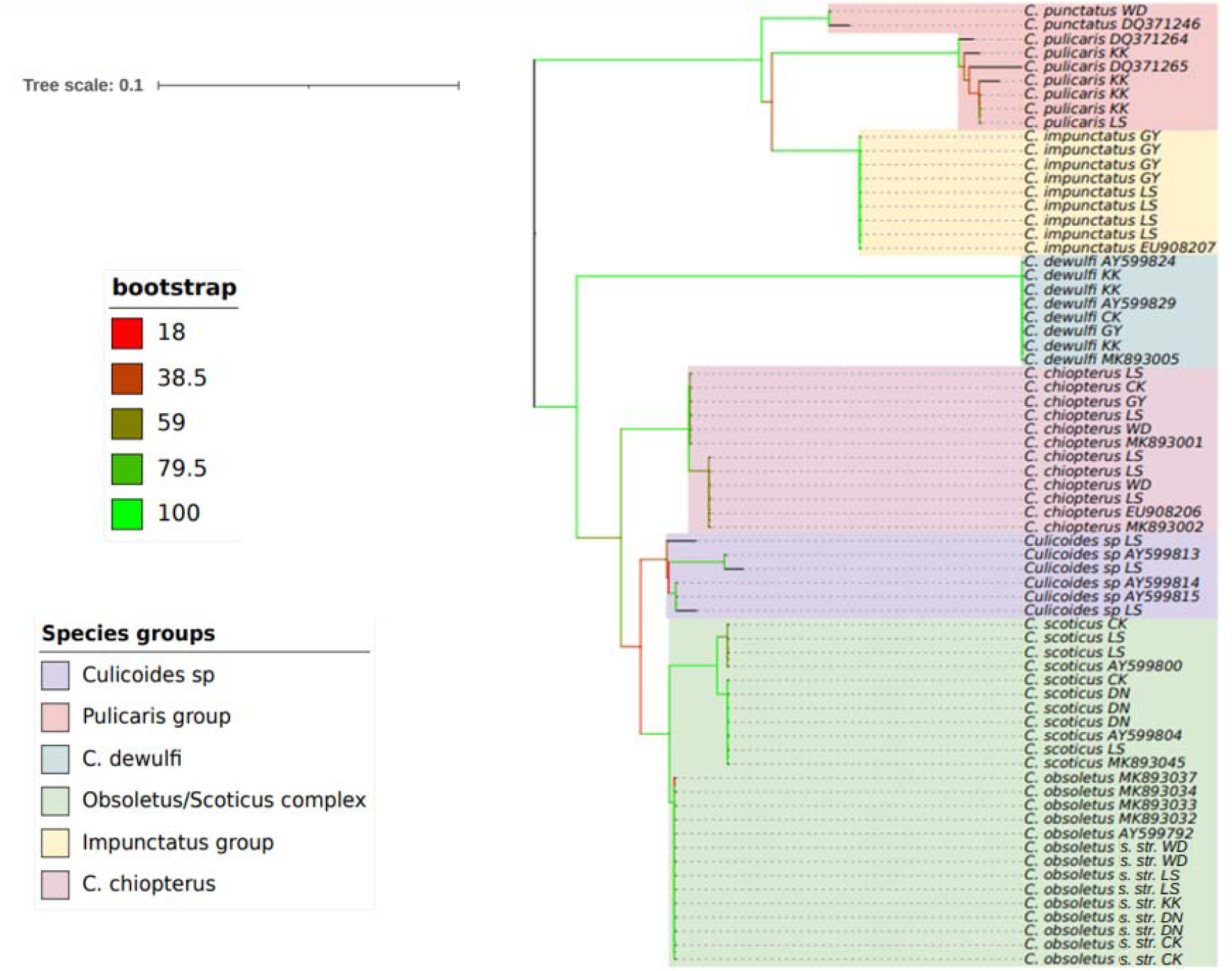
Neighbour-joining phylogenetic tree generated from an alignment of identified Irish ITS 2 *Culicoides* sequences and reference sequences obtained from GenBank truncated to a final sequence length of 218 bp. Labels on the right side indicate the species name and county code (WD – Waterford; KK – Kilkenny; CK – Cork; DN – Dublin; LS – Laois and GY – Galway).

The analysis of the CO1 and ITS2 phylogenetic trees revealed distinct groupings. According to the CO1 phylogenetic tree, three species groups were identified: the *pulicaris* group, the *impunctatus* group, and the *obsoletus*/scoticus complex, as well as the species *C. dewulfi*, and *C. chiopterus*. In contrast, the ITS2 phylogenetic tree distinguished between three species groups: the *pulicaris* group, the *impunctatus* group, and the *obsoletus*/scoticus complex, and in addition, an unidentified *Culicoides* species, *C. dewulfi*, and *C. chiopterus*. In both trees, most of the branches within clades had high bootstrap support indicating strong confidence in these evolutionary relationships. However, some branches that connect the major groups with lower bootstrap values, reflect a degree of uncertainty.

One of the most notable findings was the well-supported clade of the *obsoletus*/scoticus complex. Both, the CO1 tree and the ITS tree indicate a very close genetic relationship between *C. obsoletus s. str.* and *C. scoticus* (Figs 3 and 4) most likely reflecting this species complex. In contrast, *C. dewulfi* forms a distinct branch, separate from the other groups in both trees, representing a separate evolutionary lineage. The tight clustering of *C. dewulfi* sequences within this branch suggests a clear genetic identity, distinct from the other species. *C. chiopterus* forms a clade closely related to the *obsoletus/scoticus* complex in the CO1 phylogenetic tree but is more distantly separated in the ITS tree. The *impunctatus* group also forms a distinct clade that is separate from but closely related to the *pulicaris* group, suggesting a similar evolutionary lineage. The clustering of multiple sequences of *C. impunctatus* (Figs 3 and 4) within this clade indicates internal diversification within the species. Interestingly, the unidentified *Culicoides* species formed a distinct branch positioned between the *obsoletus*/scoticus complex and *C. chiopterus*.

### Population genetic structure of Irish Culicoides obsoletus

Analysis of the mtDNA (CO1) sequences revealed 17 haplotypes, with six haplotypes, Hap_1, Hap_2, Hap_3, Hap_4, Hap_5, and Hap_6, being the predominant haplotypes in the Irish population of *C. obsoletus s. str.* (Fig 5). Hap_1, which was the central and most prevalent haplotype in this network, was found in all counties considered in the analysis. Hap_6, Hap_3, and Hap_4 were also significant nodes connected to several other haplotypes, indicating that they are also important intermediaries in the network. Peripheral Haplotypes (Hap_5, Hap _ 8, Hap _ 9, Hap _ 10, Hap _ 11, Hap _ 12, and Hap _ 14) were more isolated and showed more specific geographic distributions.

**Figure 5:**
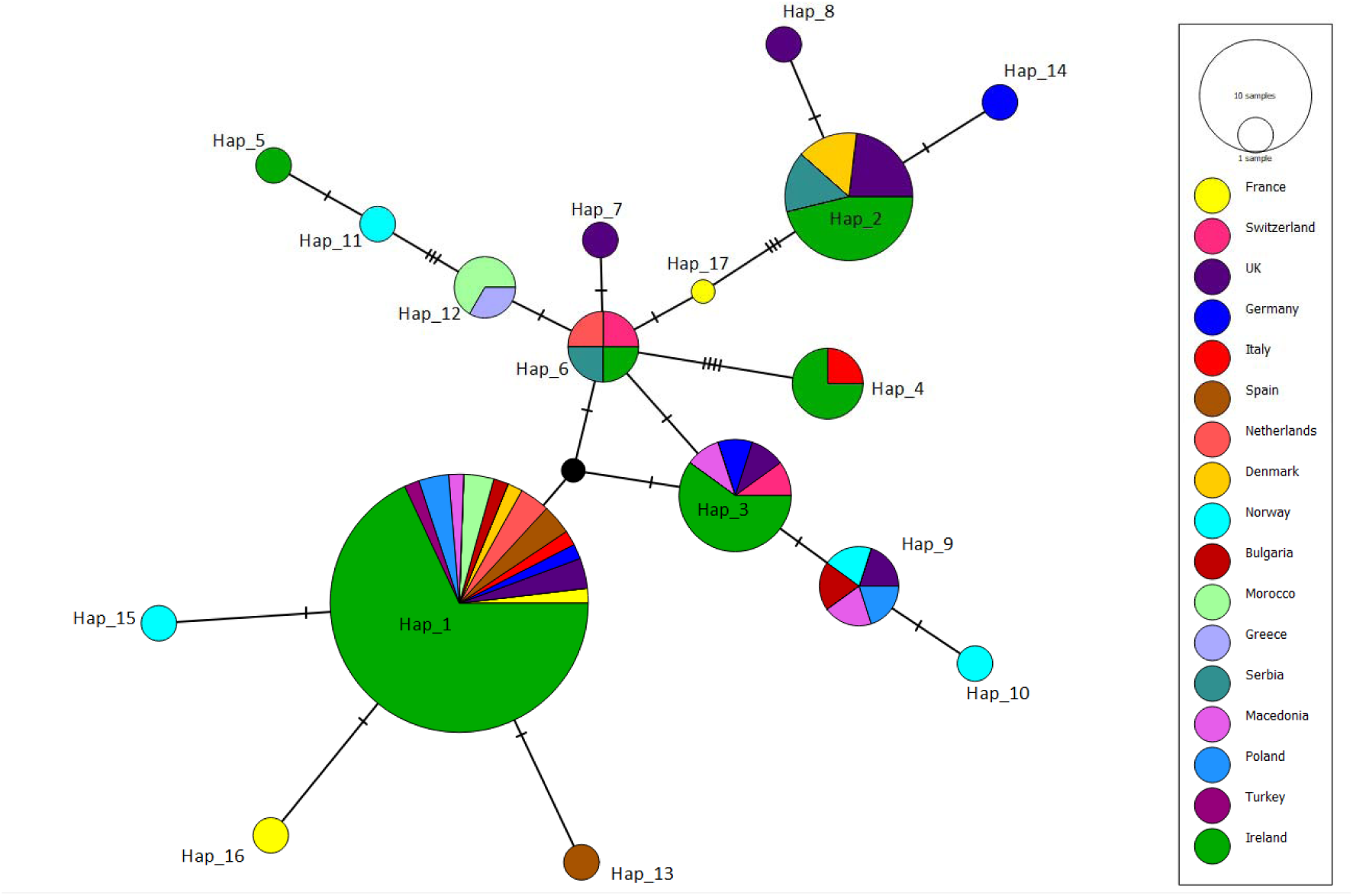
Median-Joining Network Diagram of *C. obsoletus s. str*. CO1 haplotypes from Ireland generated in this study, compared to reference CO1 sequences from other countries obtained from GenBank, truncated to 528 bp.

Analysis of the ITS2 sequences revealed just one haplotype (Hap_5) in Ireland compared to 12 haplotypes reported elsewhere (Fig 6). The haplotype Hap_5 was exclusively found in Ireland and has also been detected in Continental Europe, including France and Italy. It is closely related to other haplotypes (Hap_6, Hap_8, Hap_9, Hap_10, Hap_11, and Hap_12). In contrast, Hap_1, Hap_2, and Hap_3, found in Germany, had more mutation steps. In both network diagrams, the black node can be visualized, indicating a potential connection between the haplotypes.

**Figure 6:**
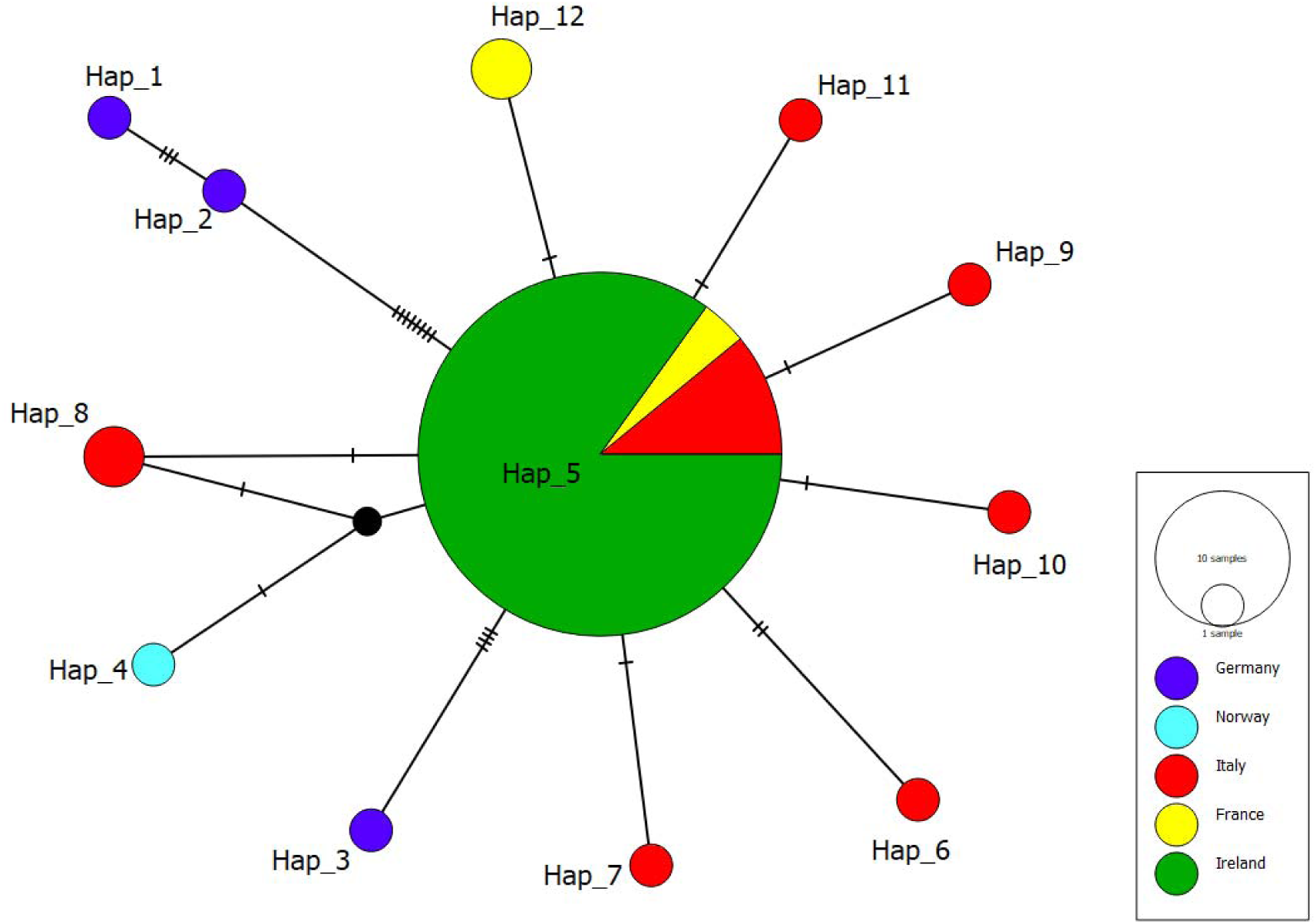
Median-Joining Network Diagram of *C. obsoletus s. str*. ITS2 haplotypes from Ireland generated in this study compared to reference ITS2 sequences from other countries obtained from GenBank, truncated to 246 bp.

## Discussion

*Culicoides* species, such as *C. obsoletus s. str.*, *C. scoticus*, *C. dewulfi*, *C. chiopterus*, *C. pulicaris, C. punctatus and C. impunctatus* have an extensive distribution across the Palearctic region, rendering them among the most widely dispersed species within this geographical area. The accurate delimitation of the status of these vector species, along with an in-depth understanding of their population dynamics and evolutionary genetics, is essential for biosecurity measures and the mitigation of *Culicoides*-borne viruses. Different molecular markers, including nuclear ribosomal DNA - (16s rDNA, 28s rDNA, ITS 1, and ITS 2), and mitochondrial DNA - (CO1, and cytochrome b oxidase (Cyt b), have been utilised to distinguish *Culicoides* species [2, 24]. The CO1 barcode is the most widely used barcode for identifying *Culicoides* species [45–49]. However, the use of this barcode can be challenging [50, 51] because of high intraspecific and low interspecific variability, leading to similar genetic distance and low genetic differentiation between species [52] Therefore, multi-locus approaches are often recommended to enhance species detection and provide accurate species composition data [29]. The ITS regions flanking the 5.8S ribosomal RNA gene evolve rapidly and are a reliable tool for species identification because sequence similarity in these regions is greater within species than between species.

In this study, both barcoding regions were used to identify common *Culicoides* species in Ireland, including species within the subgenus *Avaritia* (*C. obsoletus s. str.*, *C. scoticus*, *C. chiopterus*, and *C. dewulfi*) and the subgenus *Culicoides s. str.* (*C. impunctatus*, *C. pulicaris* and *C. punctatus*). Based on our results, ITS 2 barcode sequences were more reliable than CO1 sequences for species identification. This was particularly evident for closely related species within the *C. obsoletus* complex, where ITS 2 provided clearer differentiation between *C. scoticus* and *C. obsoletus s. str.* Similar findings have been reported in studies on species complexes in the mosquito genera *Culex* and *Anopheles* [53–55].

The diversity of *Culicoides* species detected in our study reflects the species diversity and habitat-specific distribution reported in similar studies from other geographical regions e.g. in Algeria [56]. The widespread presence of *C. obsoletus s. str.* at all trapping sites in significant numbers suggests it is a widely distributed species, possibly indicating a generalist habitat preference or its ability to thrive under diverse environmental conditions. Conversely, other species are not as widely distributed as *C. obsoletus s. str.*, may have more specialised ecological niches. *Culicoides pulicaris* and *C. punctatus* are mainly breeding in soil with high moisture levels and high organic content [27] while *C. scoticus*, *C. chiopterus*, and *C. dewulfi* [57] are often associated with livestock habitats and are known to breed in silage or manure.

*Culicoides impunctatus* was only observed at three locations: site 2 (Laois), site 4 (Galway), and site 5 (Waterford). This species typically prefers acidic, boggy grounds, areas associated with pastoral activities, elevated terrains, and regions with high sheep density [58, 59]. Known for its preference for bird hosts, *C. impunctatus* has also been recorded in ornithophilic habitats such as nest boxes [27, 60–63]. The three species classified as “*Culicoides sp*.” were recorded exclusively at site 2 in Laois. Additional molecular data from sequencing other gene markers or whole-genome sequencing, along with detailed morphological examination through specimen dissection, are needed to fully describe these species.

### Phylogenetic analysis

Phylogenetic analysis based on CO1 is considered more effective for detecting ancient hybridization in molecular phylogeny, while ITS is more reliable for resolving evolutionary issues in recently diverged taxa or cryptic species. Although the mtDNA CO1 region has been extensively used to confirm relationships within *Culicoides* [2, 23, 49], some studies have found it insufficient for analysing the closely related species within the subgenus *Avaritia*. In these cases, the use of ITS alone or in combination with CO1 has successfully clarified species relationships within this subgenus [24]. In our study, phylogenetic analysis of both the CO1 and ITS gene regions enabled the classification of the *Culicoides* species. Previous studies have shown that trimming ITS1 and ITS2 sequence alignments in *Culicoides* can result in sequences of varying lengths due to numerous indels (nucleotide insertions/deletions) in these regions as has also been reported in mosquitoes [53, 64].

Matsumoto et al. [65] observed that indel regions within ITS1 sequences divide some *Culicoides* species into two types: those with long or short ITS1 regions. This issue was overcome by analysing short ITS1 and ITS2 fragments separately after excluding indels, then concatenating them to create phylogenetic trees [24]. This approach was also employed in the present study resulting in the successful classification of the different *Culicoides* species and species groups.

Although the different species in the *obsoletus/scoticus* complex can be reliably identified based on examination of their genitalia [21, 24], adult females are challenging to distinguish morphologically due to their poorly defined wing patterns. Recent molecular characterization studies have identified nine species in the *obsoletus* group, including *C. obsoletus s. str.*, *C. scoticus*, *C. montanus* Shakirzyanova, 1962, *C. gornostavae* Mirzaeva, 1984, *C. abchazicus* Dzhafarov, 1964, and *C. filicinus* Gornostaeva & Gachegova, 1972 in the Palearctic region, and *C. alachua* Jamnback & Wirth, 1963 and *C. sanguisuga* Coquillett, 1901 in the Nearctic region and reserved the term “Obsoletus complex” primarily for *C. obsoletus*, *C. scoticus*, and *C. montanus* based on adult morphology and molecular phylogenetic analysis [2, 23, 24, 66]. During the earlier Irish surveys which were chiefly based on morphological identification, McCarthy et al. [20] used the term “Obsoletus complex” to refer to members of the subgenus *Avaritia* including *C. dewulfi*, *C. obsoletus s. lat.*, and *C. chiopterus*. Additionally, due to insufficient morphological data to distinguish female *C. obsoletus* from *C. scoticus*, these species were grouped together [21]. Using molecular tools in the present study it was possible to distinguish between *C. obsoletus s. str.* and *C. scoticus* which grouped tightly together in the phylogenetic tree (Figs 3 and 4). This close relationship suggests a high likelihood of recent divergence or ongoing gene flow between these species, resulting from potential incomplete speciation or hybridisation. A potential third member of the o*bsoletus*/scoticus complex, ‘*Culicoides* sp.’, which closely resembled specimens reported in Italy [39], was also included in the same monophyletic clade as *C. obsoletus* and *C. scoticus* (Fig. 4). These unidentified or unclassified *Culicoides* species may represent new or less-studied lineages or different variants of either the *obsoletus/scoticus* complex or of *C. chiopterus*. The close clustering of *C. chiopterus* with the *obsoletus/scoticus* complex in the CO1 tree, compared to the ITS tree, suggests that the ITS region provides stronger evolutionary differentiation between *C. chiopterus* and related species. In contrast, the CO1 gene indicates a closer evolutionary relationship between *C. chiopterus* and the *obsoletus/scoticus* complex. Phylogenetic analysis of both the CO1 and ITS2 gene regions also indicated that *C. dewulfi*, previously classified within the Obsoletus group, actually forms a distinct, monophyletic clade. This finding aligns with other studies suggesting that *C. dewulfi* only shows 86% similarity with *C. scoticus* and *C. obsoletus s. str.,* should be regarded as a distinct, monotypic entity within the subgenus *Avaritia* excluding this species from the *obsoletus*/scoticus complex. [25, 39, 45, 67].

Within the subgenus *Culicoides s. str.*, the *impunctatus* and *pulicaris* groups are well-supported clades, each characterized by high-confidence bootstrap values, indicating strong statistical support for their evolutionary relationships. The *pulicaris* group comprises 14 distinct taxa, including *C. pulicaris*, *C. lupicaris*, *C. impunctatus*, *C. punctatus*, *C. grisescens* Edwards, 1939, *C. newsteadi* Austen, 1921, *C. flavipulicaris* Dzhafarov, 1964, *C. fagineus* Edwards, 1939, *C. subfagineus* Delécolle & Ortega, 1999, *C. bysta* n. sp., *C. paradoxalis* Ramilo and Delécolle, 2013, *C. boyi* Nielsen, Kristensen & Pape, 2015, *C. selandicus* Nielsen, Kristensen and Pape, 2015, and *C. kalix* Nielsen, Kristensen and Pape, 2015, [49]. However, a study in Northern Ireland found that the *pulicaris* group in that region included only *C. pulicaris*, *C. punctatus*, and *C. newsteadi*. In that study, *C. impunctatus* and *C. grisescens* were grouped separately into an *impunctatus* group [59].

### Culicoides obsoletus network analysis

*Culicoides obsoletus s. str.*, the main vector species responsible for transmitting BTV and SBV to wild and domestic ruminants in the western Palearctic region [3, 68–71], was the most abundant species identified in this study, accounting for more than half of the *Culicoides* specimens analysed. The dominance of this species has also been reported in other studies in Europe and is attributed to its generalist nature, i.e. its ability to tolerate diverse eco-climatic conditions, and its capacity to breed in a wide range of habitats including manure in indoor locations [72, 73]. The latter is a likely reason for its significant role as a vector of livestock pathogens. Therefore, accurate delineation and understanding of its population genetic structure are important for mitigating vector-borne diseases.

The population genetic analysis of *C. obsoletus s. str.* revealed greater intraspecific variation in the CO1 region, with six haplotypes (Hap_1, Hap_2, Hap_3, Hap_4, Hap_5 and Hap_6), observed within the same *C. obsoletus s. str.* population, underscoring a wider barcoding gap in mtDNA. Conversely, just one ITS haplotype of *C. obsoletus s. str.* was revealed in Ireland and a relatively small number elsewhere, suggesting that the locus serves as a more reliable barcoding region. This has also been observed in mosquitoes where the barcoding gap for ITS ranged from 0.042 to 0.193, while for mtDNA COII, it ranged from 0.033 to 0.047 suggesting that ITS provides a more accurate molecular identification of *Anopheles* species (55).

Haplotypes characterised in the present study were compared with published sequences reported from the UK, Continental Europe, Morocco, and Turkey. The central position and widespread distribution of CO1 Hap_1 (Fig 5) and ITS Hap_5 (Fig 6) in the Median-Joining Network Diagram suggest that these may represent key ancestral haplotypes, with the other haplotypes reflecting various branches of genetic divergence. Adult *Culicoides* typically have relatively low self-propelled flight speeds compared to other insect vectors, generally covering distances of 200 to 300 meters during their lifetime [74]. However, they can travel several kilometres through passive wind dispersal, which significantly aids their colonization and spread across countries [1]. Shared haplotypes with Ireland indicate potential incursion pathways for this vector species into Ireland via passive wind dispersal. These findings support the hypothesis that previous SBV cases in Ireland [15, 22] were caused by infected *Culicoides* being carried by easterly winds leading to their colonisation and expansion in Ireland. These findings demonstrate that knowledge of the haplotype diversity in a country can be critical for understanding the incursion pathways and spread of vector-borne pathogens.

## Conclusion

This study represents the most comprehensive molecular characterisation of *Culicoides* biting midges in Ireland, in contrast to previous surveys which chiefly relied on morphological data for identification. Both barcoding regions, CO1 and ITS revealed five biting midge species within the subgenus *Avaritia* and three species in the subgenus *Culicoides*. DNA barcoding has also confirmed the presence of potential vectors of BTV and SBV in Ireland previously reported from morphological characterisation. Given the importance of the livestock sector to the economy, this is important information for understanding the potential spread of future *Culicoides*-borne pathogens. Despite their significance, the historical molecular evolutionary pathways of *Culicoides* species in Ireland remain largely unknown. However, understanding the introduction and dissemination pathways is crucial for predicting and preventing future outbreaks of *Culicoides*-borne diseases. It has been hypothesised that the current populations of *Culicoides* vector species in Ireland may have been introduced from the UK or Continental Europe. Our CO1 haplotype analysis of *C. obsoletus* elucidates various gene flow patterns, suggesting the likely introduction of midges from Europe, and possibly from North Africa and West Asia. Further research into the evolutionary history and genetic diversity of *Culicoides* populations in Ireland could yield valuable insights into their movement patterns, adaptation mechanisms, and potential for virus transmission informing biosecurity strategies and public health interventions. Moreover, the integration of molecular characterisation in *Culicoides* surveillance will help to advance research into species-specific host preference, vector competence, and preferred larval breeding sites.

## Abbreviations

BTV: bluetongue virus
SBV: Schmallenberg virus
EHDV: Epizootic Haemorrhagic Disease Virus
AHS: African horse sickness virus
CO1: cytochrome c oxidase subunit 1
ITS: internal transcriber spacer
OVI: Onderstepoort Veterinary Institute
UV: ultraviolet
NetVec Ireland: Network of Insect Vectors Ireland
BLAST: Basic Local Alignment Search Tool
MEGA: Molecular Evolutionary Genetics Analysis
iTOL: Interactive Tree of Life
PopART: Population Analysis with Reticulate Trees

## Notes

### Competing Interest Statement

The authors have declared no competing interest.

